# Mate choice in the brain: Species differ in how male traits ‘turn on’ gene expression in female brains

**DOI:** 10.1101/2023.09.15.557962

**Authors:** Jason Keagy, Hans A. Hofmann, Janette W. Boughman

## Abstract

Mate choice plays a fundamental role in speciation, yet we know little about the molecular mechanisms that underpin this crucial decision-making process. Female stickleback fish differentially adapted to limnetic and benthic habitats and considered members of reproductively isolated species use different male traits to evaluate prospective partners and reject heterospecific males. Here, we integrate behavioral data from a mate choice experiment involving benthic and limnetic fish with gene expression data from the brains of females making these mate choice decisions. We find substantial gene expression variation between limnetic and benthic species, regardless of behavioral context, suggesting general divergence in gene expression patterns in female brains, in accordance with their genetic differentiation. Intriguingly, female gene co-expression modules covary with male display traits but in opposing directions for sympatric populations of the two species, suggesting male displays elicit a genomic response that reflects known differences in female preferences which serve to isolate the species. Furthermore, our analysis confirms the role of numerous candidate genes previously implicated in female mate choice decision-making in other teleost species, suggesting that these cognitive molecular processes are, in part, evolutionarily conserved. Taken together, our study adds important new insights to our understanding of the molecular processes underlying female decision-making that maintain isolation between diverging species.

## Introduction

Choosing a mate is a key fitness decision (Andersson 1994) often crucial to speciation (Panhuis et al. 2001; Servedio and Boughman 2017; Mendelson and Safran 2021). Much research has sought to identify the male displays females assess when making mate choice decisions (Jennions and Petrie 2007; Rosenthal and Ryan 2022) as well as understand the evolution and divergence of female preferences (Ritchie 2007; Kraaijeveld et al. 2011; Servedio and Boughman 2017). However, we know very little about the cognitive and molecular mechanisms that underpin such decision-making or how these mechanisms may vary across species and depend on ecological factors (DeAngelis and Hofmann 2020; Ryan 2021). Comparative transcriptomics, where gene expression profiles are systematically analyzed across different populations, species, and environments, have already provided important insights into the evolution of mechanisms underlying various complex behaviors, such as learned vocalizations (Pfenning et al. 2014), mating systems (Renn et al. 2018; Young et al. 2019), and cooperation (Young et al. 2022). Recent transcriptomic studies have also identified some genes that appear to be involved in female mate choice (Cummings et al. 2008; Bloch et al. 2018). Yet whether and how the neuromolecular mechanisms reflecting these decision-making processes change as female preferences diverge in the process of speciation has not been examined, nor do we know how variation in gene expression contributes to reproductive isolation. Given the importance of sexual selection to both evolution and speciation, this is a critical gap. To answer these questions, we urgently need studies that examine gene expression differences in female brains as they choose among conspecific and heterospecific males.

We ask here whether and how gene expression in female brains varies among closely-related species, focusing on transcriptomic responses during courtship as females respond to variation among males in display traits known to be important in sexual isolation. We study threespine stickleback fish (*Gasterosteus aculeatus*). The limnetic-benthic threespine stickleback species pairs show parallel phenotypic evolution and parallel speciation; each species has evolved independently in multiple lakes, showing repeated and substantially parallel divergence from ancestral marine fish (Taylor and McPhail 2000; McKinnon et al. 2004). Both sexual selection and natural selection contribute to speciation (McPhail 1994; Rundle et al. 2000; Boughman 2001; McKinnon et al. 2004; Keagy et al. 2016). Limnetic and benthic species experience strong sexual isolation, but mate freely with their own species whether from their own or another lake (Rundle et al. 2000). Moreover, prior research reveals which traits females use to choose mates both within and between species; the species have diverged in female preferences for nuptial color, odor, courtship behavior, body shape and size; and females reject heterospecific males based on differences in these traits (Nagel and Schluter 1998; Boughman 2001; Kozak et al. 2009; Conte and Schluter 2013; Head et al. 2013; Mobley et al. 2016).

In our study, limnetic and benthic female stickleback were collected from two lakes where the limnetic-benthic divergence is thought to be evolutionarily independent. Once in the lab, females were courted either by a conspecific or heterospecific male from their lake or were placed with a conspecific female from their lake as a social control (Fig. 1A). We quantified male morphological and behavioral display traits, female courtship behaviors, and female preference. We generated whole brain transcriptomes for females to capture the decision-making process. We then evaluated the neural transcriptomes of females, including their relationship with behavioral and morphological data from the behavioral experiment.

**Figure 1.**
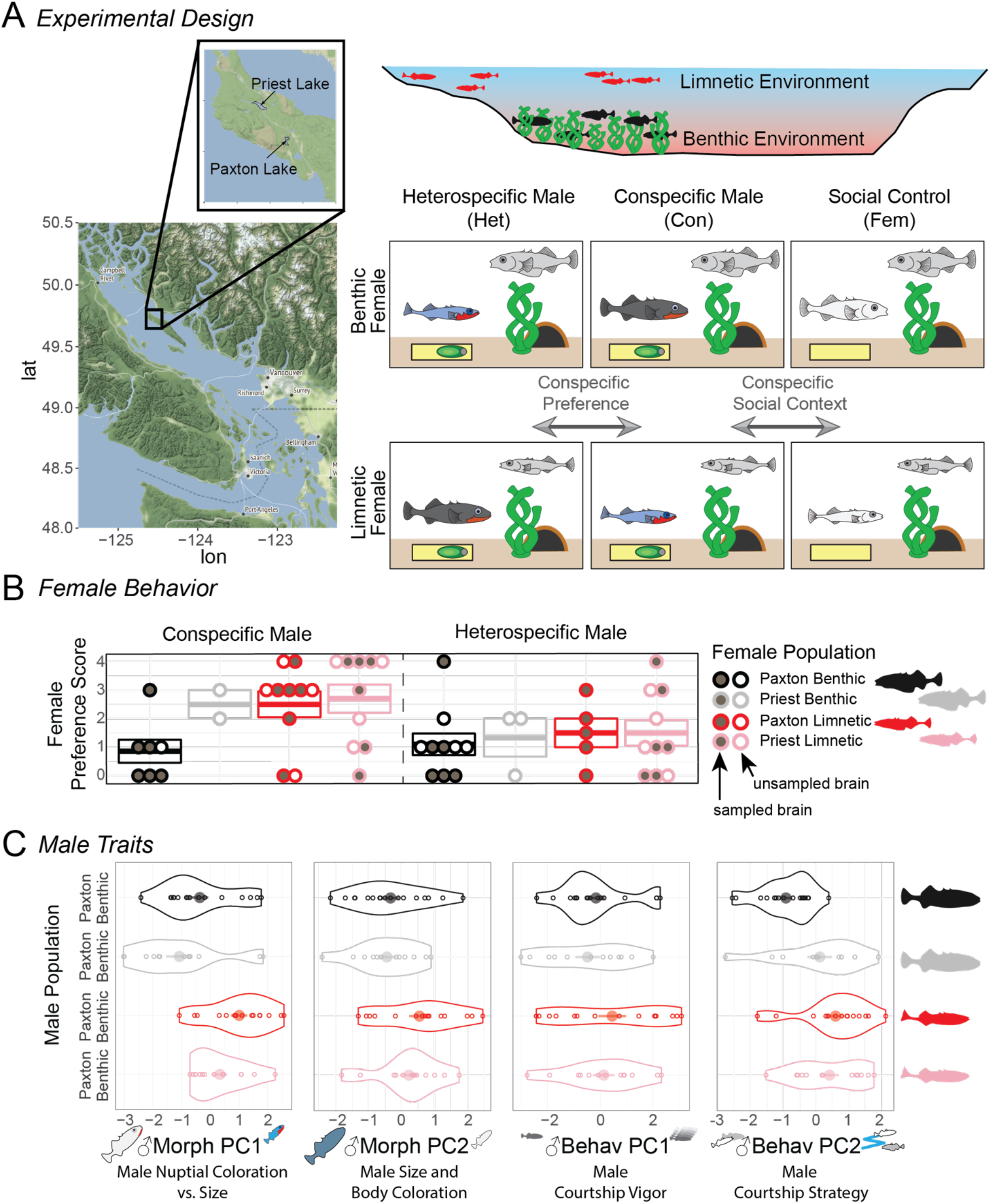
Study Overview. (A) Experimental design. Male and female stickleback fish were sampled from two lakes on Texada Island, British Columbia and brought into the lab. Females (right top corner of each panel) were exposed to three treatments before they were euthanized and brains removed: heterospecific male courtship, conspecific male courtship, or a conspecific female from their home tank. **(B) Female behavior.** Females showed stronger preference for conspecific males than heterospecific males, especially if the female was limnetic. Boxes indicate means ± SE. Each circle is an individual female. Circles that are filled in indicate females who were sampled for RNAseq. **(C) Male traits.** Five male morphological traits and five male behavioral traits were collapsed into two principal components each to summarize male variation. Male morphological PCs differentiated limnetic from benthic fish (Welch two-sample t-test: Mmorph-PC1: t_52.0_ = –4.13, P = 0.00013, Mmorph-PC2: t_54.5_ = –2.76, P = 0.0078). The first male behavioral PC (courtship vigor) did not distinguish species (t_52.06_ = –1.39, P = 0.17), but the second one, which describes well known species differences in courtship strategy, did (t_54.73_ = – 3.49, P = 0.00095).

Given that female limnetic and benthic stickleback have strong conspecific preference resulting in reproductive isolation (Rundle and Schluter 1998; Rundle et al. 2000; Kozak et al. 2009) we make several predictions. First, we predict females will show different gene expression patterns in their brains depending on the social context (conspecific courtship, heterospecific courtship, or social control) as they will not only be experiencing different stimuli, but also making different decisions with respect to reproduction. These gene expression patterns may be similar between benthic and limnetic populations, in other words, dependent on conserved molecular pathways. Alternatively, because we already know benthic and limnetic females have divergent female preferences and are influenced by different male traits (Nagel and Schluter 1998; Boughman 2001; Kozak et al. 2009; Conte and Schluter 2013; Head et al. 2013; Mobley et al. 2016), there might be important differences between the species in which genes are differentially expressed between treatments. Second, we predict that gene expression in female brains will respond to variation in male displays, and responses to male displays with a known role in sexual isolation will be in opposite directions for the two species. What we envision here is that gene expression will increase (or decrease) when the display is more attractive and the opposite will occur when the display is disliked. Because females of the two species either accept or reject males based on these display traits, the gene expression response should be opposing. The absolute direction of response for each species is difficult to predict because gene expression may ultimately lead to activation or inhibition of behavior.

Some of the strongest evidence linking female mating decisions to male displays and reproductive isolation in a neuromolecular manner would be to find gene expression patterns that are associated with *both* female choice behavior and male display in opposite directions for the two species.

## Results and Discussion

### Females preferred conspecifics males and males differed in species-specific ways

Consistent with earlier work (Rundle et al. 2000; Kozak et al. 2009; Lackey and Boughman 2014), females displayed stronger preference for conspecific males than heterospecific males (ANOVA: F_1,49_ = 11.11, P = 0.016, Fig. 1B). This was especially true of limnetic females (ANOVA: F_1,36_ = 16.90, P = 0.0062). Moreover, male displays differed between the species (Fig. 1C). We derived two principal components each for variation in female behavior (Fbehav), male behavior (Mbehav), and male morphology (Mmorph) to facilitate the integration of transcriptome data with female mating behavior and the male traits she experienced (see below, Supplementary Fig. 1).

### Gene expression quantification across populations and species

Very few of the Priest benthic females produced viable eggs, resulting in three or fewer brains per treatment for this population. Given that we were expecting large individual variation in transcriptomes, we focused our sequencing effort on the populations with larger sample sizes. After pre-processing for quality control, RNA-seq reads were mapped to the threespine stickleback (*Gasterosteus aculeatus*) coding sequences (BROAD S1 Ensembl 78 cDNA). We obtained expression information for 21,798 genes (Supplementary Table 1).

### Species show highly differentiated gene expression for individual genes

We first asked whether variation in female brain transcriptomes is concordant with previously described genetic differences among species and populations (Taylor and McPhail 1999; Taylor and McPhail 2000; Jones et al. 2012; Conte et al. 2015; Härer et al. 2021). We found that gene expression patterns were indeed highly differentiated between benthic and limnetic fish and also reflected parallel independent evolution between the two limnetic populations (Fig. 2; Supplementary Fig. 2). The extensive differences in gene expression between benthic and limnetic fish (Fig. 2; Supplementary Fig. 2) reflects their evolutionary divergence (Taylor and McPhail 1999; Taylor and McPhail 2000). Benthic and limnetic fish from the same lake had fewer differentially expressed genes, DEGs, than those from different lakes (7,443 [34% total genes] vs 8,370 [38%], adjusted p-value < 0.1, 21,796 total genes; Fig. 2B; Supplementary Table 2), possibly due to lake-specific ecology or low levels of past or contemporary gene flow (Gow et al. 2006) partly homogenizing genetic differences underlying constitutive expression patterns. Despite their independent evolution from marine ancestors, the two limnetic populations had similar overall expression patterns, as predicted by the parallel adaptation and speciation described previously (Schluter and McPhail 1992; Rundle et al. 2000; Boughman et al. 2005). Nonetheless, female brains of the Paxton and Priest limnetics did show differences in gene expression (3,642 [17%] DEGs, adjusted p-value < 0.1, 21,796 total genes; Fig. 2B; Supplementary Fig. 2C; Supplementary Table 2) and different gene expression patterns in response to specific male traits (see below).

**Figure 2.**
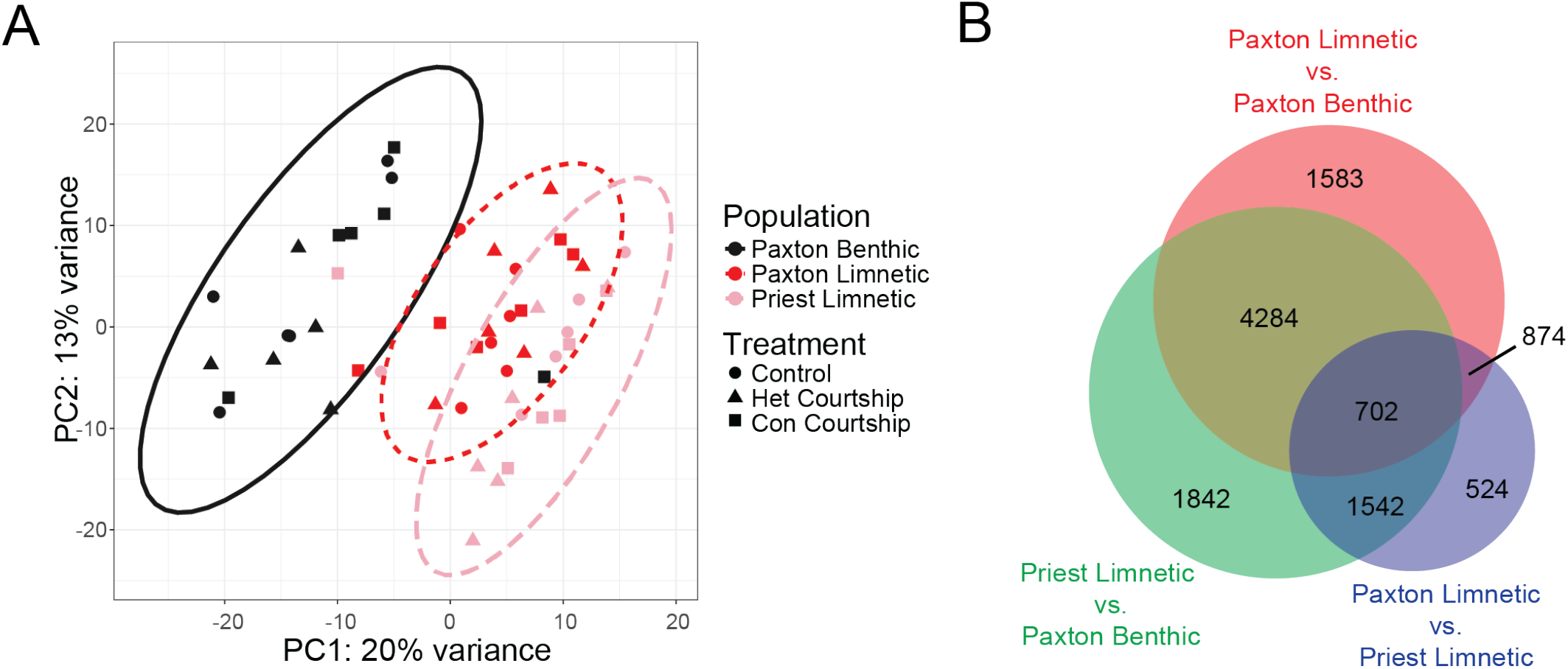
Species have highly divergent gene expression patterns. (A) Principal component analysis (PCA) of variance stabilized normalized counts of 90% most variable genes. PC1 clearly separates populations in expected ways. For example, limnetic populations overlap and are very well separated from the benthic population. The limnetic population that is most similar to the benthic population is the one from the same lake. Ovals are 95% confidence ellipses. Two samples appeared to have been swapped (Priest limnetic and Paxton benthic; they are at the center of the other’s population distribution) and are not included in further analyses. **(B) Venn diagram comparing overlap of population comparison differentially expressed genes (DEGs, FDR corrected p-value < 0.1).** Again, there were many more genes that distinguished benthic from limnetic fish than limnetic fish from different lakes.

### Magnitude of gene expression differences between treatments reflect species differences in strength of courtship responsiveness and conspecific preference

We euthanized all fish within 20 min stimulus onset to reveal gene expression during decision-making. Even with this short stimulus time, limnetic females from both populations showed numerous DEGs in comparisons between the conspecific male treatment versus either the conspecific female or heterospecific male treatments (28-52 [0.13-0.24%], adjusted p-value < 0.1, 21,796 total genes; Fig. 3; Supplementary Table 3). Benthic females, in contrast, had fewer DEGs for both of these comparisons (7, Fig. 3), consistent with prior behavioral work indicating limnetic females are more responsive to male courtship and have stronger conspecific preferences than benthic females (Boughman et al. 2005; Kozak et al. 2009). These patterns are interesting in light of earlier findings in the Panuco swordtail (*Xiphophorus nigrensis*), where females strongly prefer large courting males to small coercive males (Ryan and Rosenthal 2001). This robust behavioral preference in swordtails was reflected in the brain transcriptome: the brains of females in this choice situation have many more DEGs than females exposed to only small males or only females (Cummings et al. 2008). Recent work in another poeciliid species, the guppy (*Poecilia reticulata*), also demonstrated that females that expressed a strong preference for colorful over drab males had many more DEGs than females that did not have a preference regarding male coloration (Bloch et al. 2018). Taken together with our findings, these results point to stronger gene expression response with greater mate preference.

**Figure 3.**
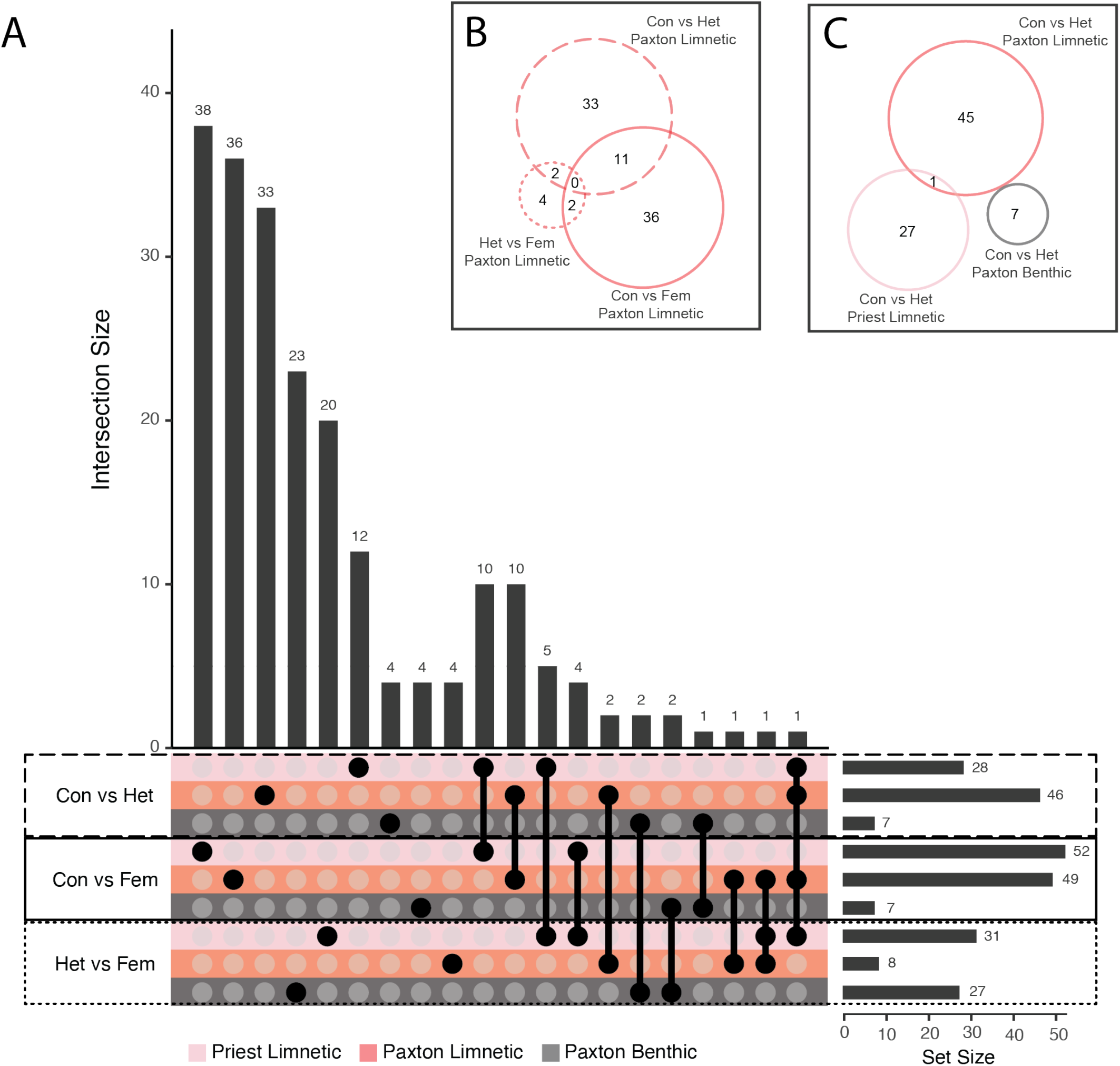
Differentially expressed genes from treatment comparisons. (A) UpSet plot showing all treatment comparisons separated by population. Solid circles connected by lines indicate the intersection between those sets – circles that are unconnected to any others indicate genes unique to that set. Generally, there is very little overlap between sets; when there is, it is between different comparisons within the same population or species. We used a significance threshold of FDR corrected P < 0.1. Traditional Venn Diagrams can be derived from the UpSet plot as seen in **(B) Paxton limnetic treatment comparisons** and **(C) Conspecific versus heterospecific comparisons.**

### Genes involved in conspecific preference are population-specific

Support that female brains are differentiating conspecific from heterospecific males, and that gene expression is involved in this decision-making, are the DEGs found by comparing the conspecific male to heterospecific male treatments (“conspecific preference”, 7-46 DEGs, [0.03-0.21%], adjusted p-value < 0.1, 21,796 total genes; Fig. 3C; Supplementary Table 3). If female gene expression was not involved in discriminating between conspecific and heterospecific males, we would expect no DEGs in this comparison. Instead, our results suggest that conspecific and heterospecific males elicit differences in expression of certain genes in female brains. Notably, however, we found virtually no overlap in DEGs between populations for conspecific preference (Fig. 3C), suggesting that different genes are involved in conspecific preference in each population. We expected substantial differences for benthic compared to limnetic populations because they have distinct mating preferences and social behavior (Vamosi 2002; Boughman et al. 2005; Kozak et al. 2009); our findings support that prediction and suggest that behavioral and gene expression differentiation go hand-in-hand. However, we expected some overlap between limnetic populations because as they evolved from marine ancestors in mating and social behavior, they did so in parallel (Vamosi 2002; Boughman et al. 2005; Kozak et al. 2009), and because of their overall similarity in gene expression discussed above. The little overlap we found for limnetic conspecific preference accords with independent evolution of the species pairs in each lake (Taylor and McPhail 2000), which appears to result in distinct molecular networks related to conspecific preference (reflected also by non-overlap of Gene Ontology (GO) terms, Supplementary Table 4). This could have been either due to the available genetic variants differing (*sensu* the mutation order hypothesis (Mani and Clarke 1997; Schluter 2009; Mendelson et al. 2014)), or because selection was less similar than thought. Although there are exciting examples of parallel gene expression changes underlying parallel evolution of complex behavioral phenotypes (Pfenning et al. 2014; Renn et al. 2018; Young et al. 2019; Jacobs et al. 2020; Young et al. 2022), it seems that parallel phenotypic evolution often rests on only partly parallel genetic mechanisms (Soria-Carrasco et al. 2014; Bolnick et al. 2018).

### Gene co-expression network analysis supports individual gene analysis results

We next used weighted gene co-expression network analysis (WGCNA (Langfelder and Horvath 2008; Langfelder and Horvath 2012)) to identify gene modules that may underpin shared biological functions (Supplementary Fig. 3, Supplementary Table 5). We identified 13 differentially expressed module eigengenes (DEMEGs) between limnetic and benthic female brains from the same lake (Fig. 4; Supplementary Table 6), similar to patterns we uncovered with the individual gene analyses. Also similarly, we found fewer DEMEGs when comparing limnetic females from different lakes (7 DEMEGs; Fig. 4; Supplementary Table 6). However, five of these seven modules also differentiate Paxton limnetic from Paxton benthic fish and so differentiate populations generally. Thus, there is very strong differentiation in module eigengene expression between species and weaker differentiation between limnetic populations, supporting our prediction of divergent gene expression between diverged species. Although we found relatively few DEMEGs between treatments (Fig. 4; Supplementary Table 6), we did identify two DEMEGs when females were courted by a conspecific versus heterospecific male; one for Paxton benthic and one for Paxton limnetic females. We suggest that these modules are involved in conspecific recognition and may play an important role in sexual isolation. We also found three DEMEGs when benthic females were courted by a conspecific male versus interacted with a female, suggesting these modules are involved in evaluating males. We were surprised that benthic females showed a strong effect here as they neither exhibited strong preference nor strongly discriminated between males at the behavioral level (Fig. 1B) and they had the fewest DEGs for these comparisons (Fig. 3; Supplementary Table 3). Module eigengene expression likely reflects female perception and decision-making rather than a final decision given the relatively short time (≤20 min) between trial start and sampling brains. We therefore examined how female module eigengene expression relates to male display trait variation and female choice behavior.

**Figure 4.**
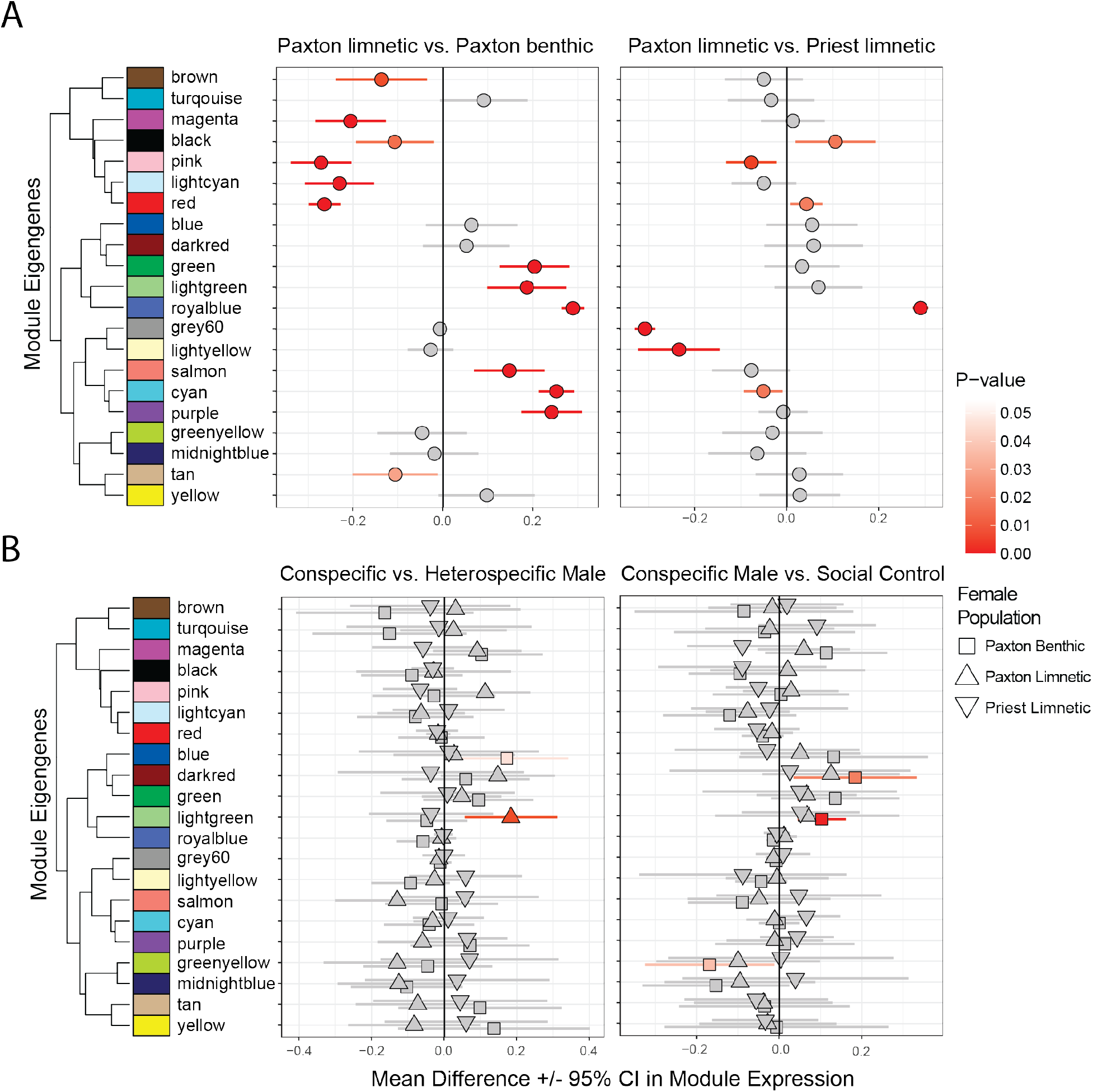
Comparisons of module expression between (A) populations and (B) treatments. Symbols and lines correspond to mean ± 95% confidence intervals for t-tests. Significant differences are indicated in color.

### Module eigengene expression in the female brain reflects individual variation in a male sexually selected trait in population-specific ways

It is well established that variation in male traits influences both current and future female mating decisions (Andersson 1994; Jennions and Petrie 2007; Tinghitella et al. 2013; Rosenthal and Ryan 2022), the specific traits that females focus on vary between populations (Endler and Houde 1995; Scordato and Safran 2014), and changes in the expression and topology of gene networks interface with neural networks to influence future behavior (Sinha et al. 2020). We therefore predicted that module eigengene expression in the female brain should vary in conjunction with male displays and female behavior, likely in a species– or population-dependent fashion. Indeed, this was true for an important male morphological trait, throat color, which varies between benthic and limnetic species, is involved in female choice and male competition, and is critical to sexual isolation (Boughman 2001; Albert et al. 2007; Boughman 2007; Lackey and Boughman 2013; Keagy et al. 2016). Variation in male throat color significantly predicted variation in the eigengene expression of two modules in a population-dependent manner (Fig. 5; Supplementary Table 7), with mean expression and slopes differing between benthics and at least one limnetic population, especially the sympatric one. Thus, the activity of brain gene co-expression modules was altered in female brains in response to this key male trait known to be subject to sexual selection and involved in sexual isolation.

**Figure 5.**
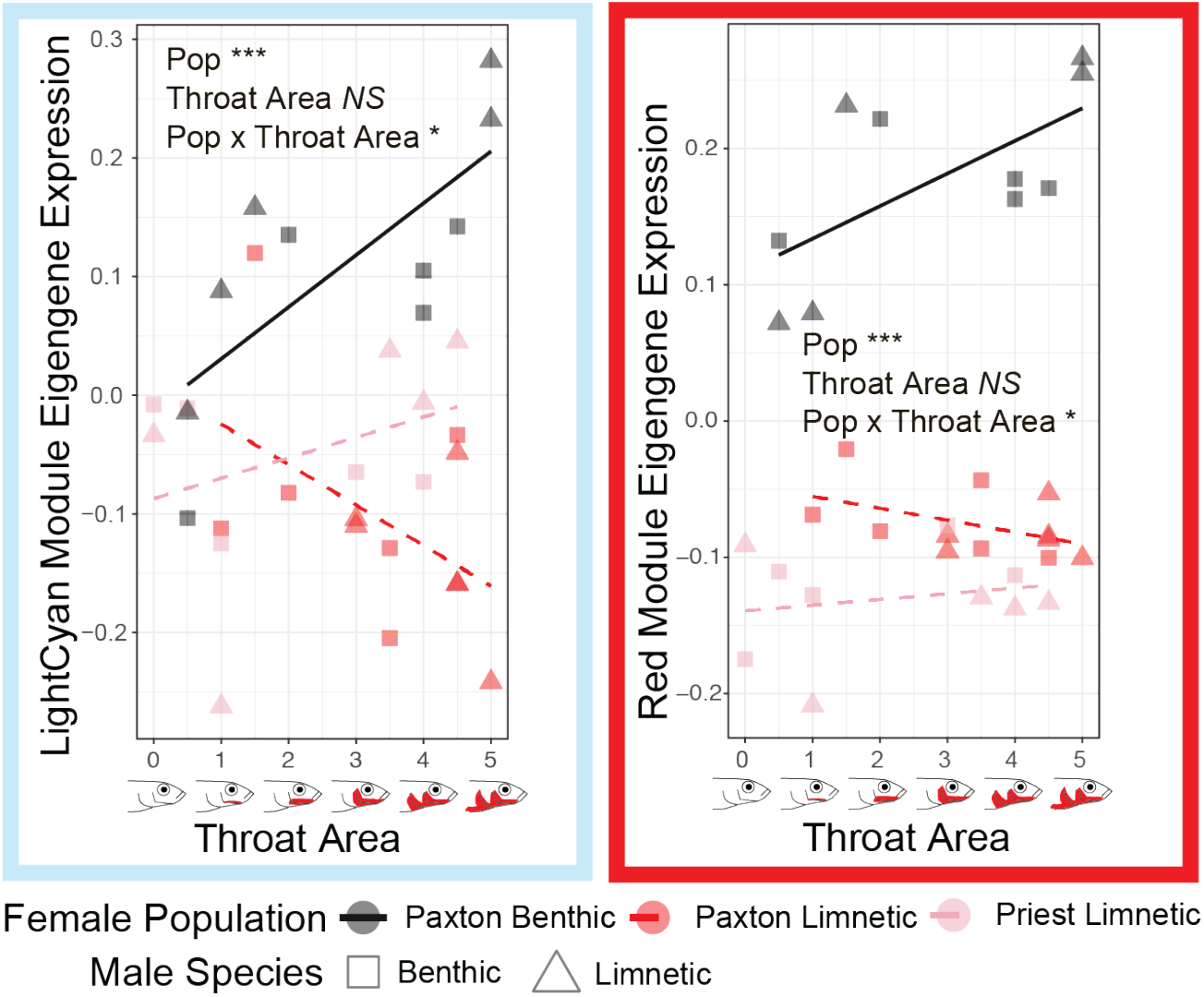
Gene co-expression modules whose expression in female brains are predicted by male throat color, but in different ways depending on population. Significance of the fixed effects of population, traits, and their interaction are indicated: *** < 0.001, ** < 0.01, * < 0.05.

Importantly, the relationship between module eigengene expression and male throat color was strongly divergent in benthic and limnetic females from the same lake, in parallel with strongly divergent preferences for this trait in these same females (Boughman 2001). This finding argues that the differential recruitment of gene networks implied by these co-expression modules underpins divergence in mating behavior and conspecific preference, thus contributing to reproductive isolation between diverging species.

Our finding that module eigengene expression responds to male throat color helps to explain the low number of DEMEGs for treatment comparisons. Based on these results, females in the same treatment would be expected to have variable expression of some modules based on the trait values of the specific males courting them. Furthermore, although limnetic males tend to have more coloration than benthic males, there is overlap in the distribution, leading to imperfect correspondence between coloration and treatment and likely obscuring the gene expression signal from treatment comparisons.

### Moving from individual traits to multivariate descriptions of male & female phenotypes

Mate choice decisions are typically based on more than a single trait (Endler and Houde 1995; Gerhardt and Brooks 2009; Oh and Shaw 2013; Ryan 2021). We therefore used the axes of variation in male behavior (Mbehav PC1 and PC2) and morphology (Mmorph PC1 and PC2) as well as female behavior (Fbehav PC1 and PC2) we previously inferred using PCA (Supplementary Fig. 1) to integrate variation in these traits with our gene expression data. We did this using linear models, testing for population and trait PC main effects as well as their interactive effects on module eigengene expression (Fig. 6; Supplementary Table 7). Significant main effects of trait PCs (with no population by trait interaction) indicate modules that respond to specific trait PCs in a consistent manner across all populations. Genes in these modules could reflect conserved patterns of gene expression during courtship. We did find this pattern with the PC describing male size and body coloration (Mmorph PC2) for 4 modules, the PC describing male courtship vigor (Mbehav PC1) for 3 modules, and the PC describing the nature of female response (Fbehav PC2) for 1 module, for a total of 5 unique modules. Several of these modules have clearly differentiated biological functions, according to GO analysis (Supplementary Table 8), giving some clues about the conserved molecular mechanisms underlying female mate choice decisions. For example, the turquoise module (Mmorph PC2 and Mbehav PC1) is involved in protein transport, metabolism and biosynthesis of various molecules, mitochondria and energy functions, and DNA/RNA processing. The brown module (Mbehav PC1) is involved in metabolism and biosynthesis, translation, and cellular stress response. The yellow module (Mmorph PC2) is involved in synaptic transmission, ion transport, and cell-cell signaling.

**Figure 6.**
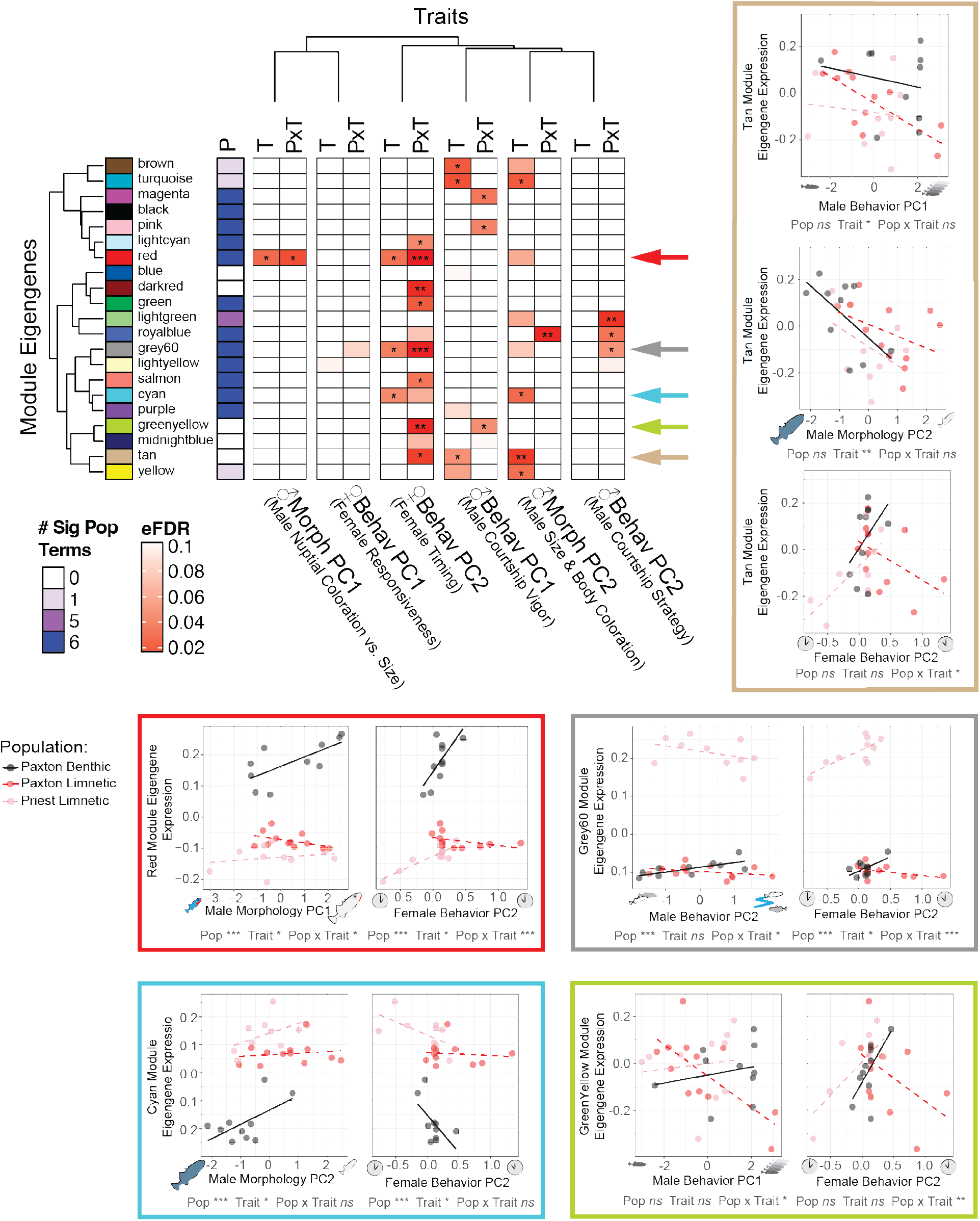
Linear models describing relationship between module eigengene expression and population, traits, and their interaction. Arrows indicate modules that respond to both a male display trait and female courtship behavior; these relationships are further explored in scatterplots. “P” = population term, “T” = trait term, “PxT” = population by trait interaction term with significance indicated: *** < 0.001, ** < 0.01, * < 0.05. To focus on the trait and trait by population interaction effects, we have reduced the population term to a single column indicating in how many of the six models it was significant (at P < 0.05).

Most importantly, however, significant population by trait interactions address our primary prediction that divergent selection on female mate preferences would result in divergent brain gene expression patterns in response to male display trait values, especially in benthic and limnetic females from the same lake. Indeed, we saw many modules with this pattern; 7 modules showed a significant male trait by female population interaction and, in all cases, limnetic and benthic females from the same lake had opposing slopes (e.g., see Fig. 6). When we shifted our focus to female behavior in response to courtship, we found a similarly large number of modules (n = 8) that showed a population by trait interaction, specifically the nature of female response (early vs. late, Fbehav-PC2). Once again, we observed that limnetic and benthic females from the same lake had opposite slopes, showing strong divergence in gene expression as predicted by their opposing trait preferences.

GO analysis of the 8 modules with a significant population by Fbehav-PC2 interaction indicates distinct biological functions being overrepresented by module genes (Supplementary Fig. 4; Supplementary Table 8), especially for the greenyellow and green modules: DNA/RNA processing, metabolic processes, and cellular stress response (greenyellow) and synaptic signaling, ion transport, cell-cell signaling, and neural development (green). Clearly something very different is happening in female brains of sympatric species of stickleback. The preponderance of genes with neural functions (green module) and those influencing future gene expression and response to stress (greenyellow module) suggests those modules are mediating activity in the brain involved in decision-making. The different expression patterns for limnetics and benthics from the same lake point to a key role for these modules in isolating the species.

Even more interesting, for five modules there was overlap where a module’s eigengene expression was associated with both male trait and female behavioral variation (Fig. 6). We interpret this to indicate that the male trait elicits a specific neurogenomic response in the female brain that, in turn, influences female choice behavior. For example, the red module showed lower eigengene expression in Paxton limnetic females when they were courted by males that they were expected to find more attractive (smaller and with brighter blue eyes and more extensive red throat coloration). These females in turn had lower eigengene expression of the red module if they showed more interest later in courtship. Paxton benthic females showed the exact opposite relationships between red module eigengene expression and male morphology and female behavior. The finding of overlap in modules showing associations with male traits and female behavior suggests these modules are important for premating isolation.

### Candidate gene expression also associated with trait values

Numerous candidate genes have previously been implicated as playing a role in female mate choice and social decision-making more generally (Weitekamp and Hofmann 2017; DeAngelis and Hofmann 2020). We therefore mined our transcriptome dataset for candidate genes representing five distinct and well-studied neuroendocrine and neuromodulatory pathways. Specifically, we focused on genes involved in 1) **gonadotropin-releasing hormone (GnRH) signaling**, due to their role in reproduction (Gore 2002): *gnrh1, gnrh2, gnrh3, gnrhr4*; 2) **nonapeptide signaling**, known for regulating affiliative behavior (Goodson 2008): *avp, avpr2, oxt, oxtr*; 3) **dopamine signaling**, which is important for motivational processing (Lerner et al. 2021): *th, th2, DRD1, drd1b, drd2a, drd2l*; 4) **prostaglandin F2 alpha (PGF2α) signaling**, as the ovarian hormone PGF2α is a well-known regulator of reproductive behavior in fishes (Kidd et al. 2013; Juntti et al. 2016): *ptgfr*; and 5) specific genes important to **synaptic plasticity** that have previously been implicated in mate choice decisions in poecillid fishes (Cummings et al. 2008; Wong et al. 2012; Bloch et al. 2018): *nlgn1, nlgn2a, nlgn2b, nlgn3a, nlgn3b, neuroligins, neuroserpin1*. We found that 20 of these 24 genes (83%) were members of 9 different gene co-expression modules (Fig. 7), with the blue (5 genes) and brown modules (4 genes) most prominently represented (recall that the brown module was correlated with Mbehav PC1, courtship vigor). We then used linear models as with the gene co-expression modules above to discover that 14 genes (58%), representing four of these pathways (all except for PGF2α signaling), showed significant differences in expression between populations (Fig. 7, Supplementary Table 9).

**Figure 7.**
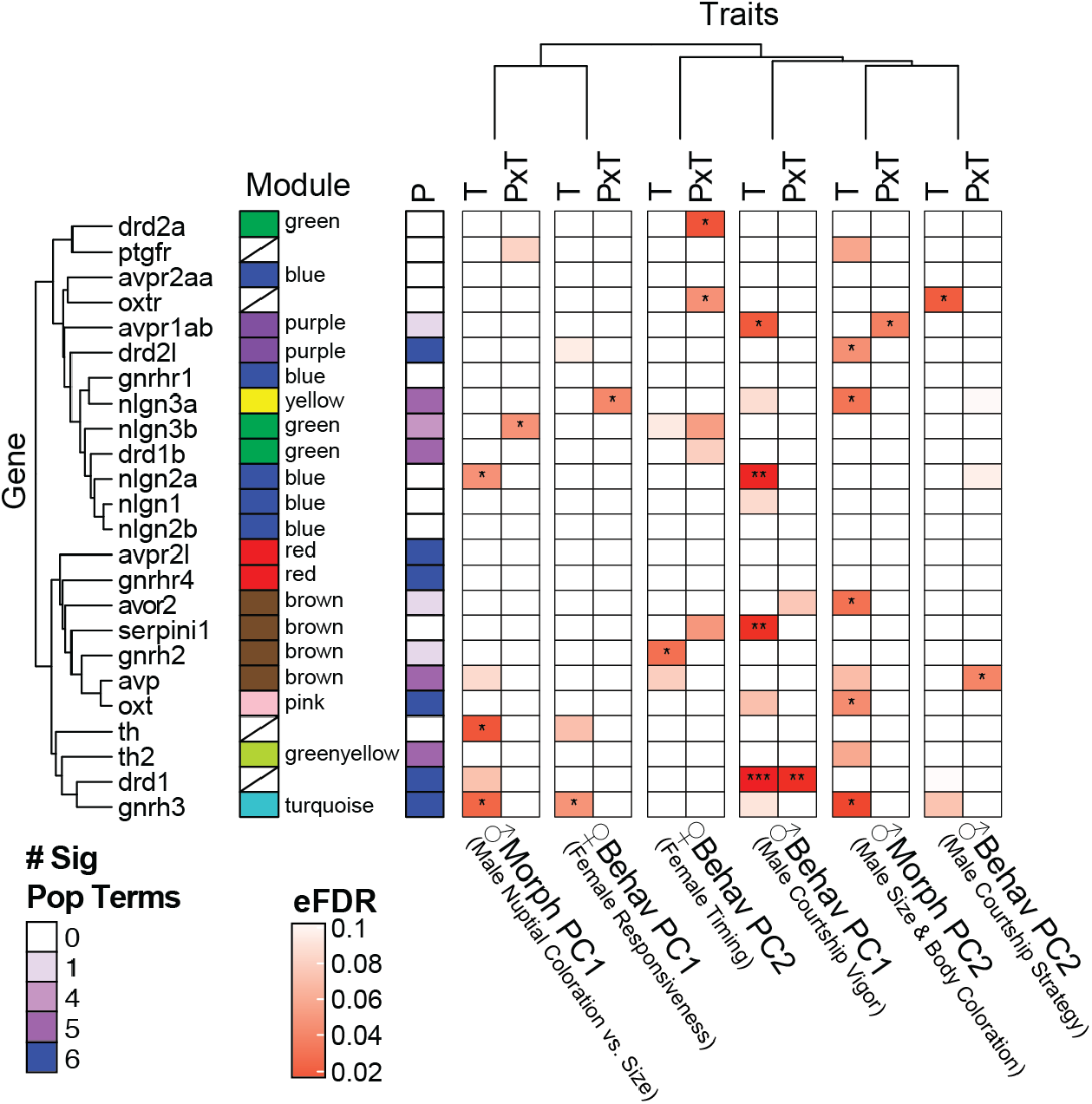
Linear models describing relationship between candidate gene expression and population, traits, and their interaction. “P” = population term, “T” = trait term, “PxT” = population by trait interaction term, with significance indicated: *** < 0.001, ** < 0.01, * < 0.05. To focus on the trait and trait by population interaction effects, we have reduced the population term to a single column indicating in how many of the six models it was significant (at P < 0.05).

When we focused on significant relationships between candidate gene expression and specific male or female traits, we discovered several intriguing associations. For example, the expression of GnRH3, which has previously been shown to gate mating preferences in medaka (Japanese rice fish, *Oryzias latipes*) (Okuyama et al. 2014), was associated with female responsiveness (Fbehav PC1), while GnRH2 expression reflected female timing (Fbehav PC2). Note that GnRH2 plays a critical role in the integration of energy homeostasis and sexual behavior in mammals and teleosts (Temple et al. 2003; Marvel et al. 2021). Looking at the behavior and morphology of the males, we found that dopaminergic and nonapeptide signaling along with synaptic plasticity in the female brain reflect male courtship vigor (Mbehav PC1), while oxytocin receptor expression is related to male courtship strategy (Mbehav PC2).

Similarly, GnRH and dopaminergic signaling along with synaptic plasticity reflect a male’s nuptial coloration (Mmorph PC1). Finally, male size and body coloration (Mmorph PC2) predicted (albeit weakly) PGF2α signaling in the female brain. Taken together, our candidate gene analysis is consistent with findings in other species and, importantly, also suggests several avenues for future research.

## Conclusion

Despite their close evolutionary relationship, we find substantial differences in gene expression for limnetic and benthic females making mate choice decisions. This differentiation is not due simply to differences in magnitude of expression in the brains of the two species, but also due to opposing patterns in response to male displays such as color and courtship behavior, as well as female choice behavior. Our most important and novel finding is that brain gene expression responds to male display traits in opposite directions for the two sympatric species, mirroring contrasting female behavioral responses to those displays known to contribute to sexual isolation as shown here and by earlier work. Male displays that trigger elevated expression of a module in female brains for one species trigger reduced expression in the other species. Thus, the divergent male displays and divergent female preferences of the limnetic and benthic species are matched by divergent gene expression in female brains in association with male display and female choice behavior variation. This supports our key prediction that not only should there be differentially expressed genes in benthic and limnetic fish, but this differential expression is driven by the diverged female preferences for diverged male display traits in a quantitative manner.

## Materials and Methods

Benthic and limnetic threespine sticklebacks, *Gasterosteus aculeatus*, were collected from two lakes on Texada Island, British Columbia (BC): Paxton (49°42’36“N 124°31’30“W) and Priest (49°44’42“N 124°33’54“W) in April 2014. Fish were transferred to Michigan State University, where they were housed in 110 L aquaria, in densities of up to 30 individuals per tank, separated by lake and species. We used a timer system that changed the light:dark cycle continuously to track natural changes in daylight in BC. Temperature was maintained at 15°C. Animals were fed defrosted brine shrimp (*Artemia* sp.) and bloodworms (*Chironomus* sp.) *ad libitum* once per day.

## Behavioral trials

We used standard methods for assessing female mate preference in stickleback fishes in which a female is courted by a single male (Nagel and Schluter 1998; Kozak et al. 2009; Tinghitella et al. 2013). Each female subject experienced one of three trial conditions: 1) courtship by a male from their lake and species (conspecific courtship), 2) courtship by a male from their lake and the opposite species (heterospecific courtship), or 3) a control in which females were with a female from her same lake and species (social condition) (Fig. 1A).

During May – August 2014, reproductive males were taken from holding tanks and placed individually in visually isolated 110-L aquaria (with a 76 x 31 cm footprint and water depth of 43 cm) with nesting materials and enticed to build a nest (see Supplementary Materials and Methods for more details).

All behavioral trials commenced between 0930 ET and 1230 ET. We verified each female’s reproductive status by gently squeezing her abdomen to confirm presence of ripe eggs in the oviduct (Head et al. 2013). For the courtship treatments, females were placed in an opaque holding container just below the water surface in the male’s nesting tank for a 5 min acclimation period. The female was then remotely released, and the behaviors of both male and female were recorded using the event recorder JWatcher 1.0 (http://www.jwatcher.ucla.edu/) by two observers. The social control treatment was done similarly except that the stimulus fish was a non-reproductive female from the subject female’s home tank.

After 20 min or spawning (whichever came first, see Supplementary Materials and Methods for more details), the female was killed by rapid cervical dissection, and the brain was immediately dissected under a microscope in a Sylgaard 184-lined petri-dish filled with a Ringer’s solution. Brains were stored overnight in RNAlater at 4°C and then transferred to –20°C until RNA extraction. Presence of viable eggs for all females was confirmed by retrieving eggs from the nest or, if no spawning had occurred, gently squeezing the female carcass.

## RNA-seq

In September 2014, brains were shipped on dry ice to The University of Texas at Austin where RNA was extracted using the Maxwell 16 LEV simplyRNA Tissue Kit, which utilizes a robot increasing consistency. At the UT Austin Genomic Sequencing and Analysis Facility each sample (all with RIN > 7.8, 8.8 ± 0.4 mean ± SD, 11 samples do not have RIN scores) was prepared for RNA-seq using Poly-A mRNA capture and given a unique barcode. A single library of all multiplexed samples was sequenced across eight lanes of an Illumina Hiseq 2500 with 2×50 PE chemistry. Sample size for each treatment for Paxton limnetic, Paxton benthic, and Priest limnetic populations was n∼6. We did not sequence any samples from the Priest benthic population because of small sample sizes. Bioinformatic analyses were carried out using the computational resources of the Texas Advanced Computing Center (TACC).

RNA-seq resulted in 34.2 ± 4.6 million reads (mean ± SD) per sample per sequencing direction (forward or reverse). Reads from all eight lanes were combined for each uniquely barcoded individual, separately for forward and reverse reads. We conducted quality control checks using *fastqc* (0.11.1). We then ran *Trimmomatic* v0.33 to remove a small amount of adapter contamination. Next, reads were aligned to the *G. aculeatus* Ensembl BROAD S1 draft genome (version 78) using *bwa* v0.7.7. *Samtools* (1.2) was used to convert sam to bam files, and then sort and index them. These sorted and indexed bam files were then passed to *bedtools* (2.23.0) to count gene transcripts. The resulting gene counts were analyzed quantitatively using *DESeq2* (Love et al. 2014) and *WGCNA* (Langfelder and Horvath 2008; Langfelder and Horvath 2012) in *R* v4.2.2 (R Core Team 2022) as described below.

## Analysis

Additional detail about analyses of female behavioral and male behavioral and morphological data are described in the Supplementary Materials and Methods.

Using the *rlog* function in the *DESeq2* library (Love et al. 2014) and a design matrix with an intercept only, we first applied a ‘regularized log’ transformation to the gene expression count matrix which was exported for the WGCNA and candidate gene analysis (Supplementary Table 10, more below). Using these normalized and variance-stabilized gene expression counts we conducted a principal components analysis on the 90% most variable genes as an initial visualization step. Two samples appeared to have been swapped and were removed from analyses (Fig. 2A). The original untransformed gene expression count matrix was then analyzed using standard methods (Love et al. 2014) with a model where gene expression counts were predicted by the independent effects of treatment and population. Subsequent contrasts were computed to answer whether there were species differences (Paxton limnetic vs. Paxton benthic), lake differences (Paxton limnetic vs. Priest limnetic), and population differences using the “normal” shrinkage option and an alpha of 0.1. Treatment difference contrasts were calculated separately for each population from a second model in which gene expression counts were predicted by the interactive effect of treatment and population. The *gprofiler2* library was used for GO analysis (Kolberg et al. 2020).

WGCNA was conducted using standard methods for signed network construction using functions in the *WGCNA* library (Langfelder and Horvath 2008; Langfelder and Horvath 2012). Briefly, we used the 90% most variable genes from the normalized and variance-stabilized gene expression count matrix (without the two samples suspected of being swapped). Additional quality control steps eliminated two samples that had the poorest read-mapping percentage.

Using the Texas Advanced Computing Center (TACC), we tried different combinations of parameters to optimize network construction rather than simply relying on defaults. The analysis presented here was done with modules created using the following parameters: maxBlockSize = 30,000, power = 9, network-type = “signed”, corType = “bicor”, maxPOutliers = 0.05, minModuleSize = 30, mergeCutHeight = 0.25, deepSplit = 2 (default), and detectCutHeight = 0.995 (default). Modules were analyzed first using t-tests to compare species (Paxton limnetic vs. Paxton benthic), lakes (Paxton limnetic vs. Priest limnetic), and treatments. Then we constructed linear models (*lm* function in the *stats* library (R Core Team 2022)) with module eigengene expression predicted by population, trait, and their interaction. We assessed statistical significance of main and interaction effects using an empirical FDR procedure (Storey and Tibshirani 2003). Briefly, significance was initially calculated using a analysis-of-variance table with type-II sums of squares from the linear model results (using the *Anova* function in the *car* library (Fox and Weisberg 2019)). Then the gene expression counts were shuffled among samples 10,000 times and new significance values calculated each time. eFDR was the proportion of times the true significance value was lower than or equal to the significance values calculated from these shuffled datasets. The *gprofiler2* library was used for GO analysis (Kolberg et al. 2020).

Candidate gene expression was also analyzed using linear models with gene expression predicted by population, trait, and their interaction and significance assessed using an empirical FDR procedure.

## Data, Materials, and Software Availability

Data and code will be uploaded to Dryad. Raw sequencing reads will be deposited in GenBank. All other study data are included in the article and/or Supplementary Material.

## Acknowledgements

Research was conducted under permits from the Ministry of the Environment, BC and approval of the Institutional Animal Care and Use Committee of Michigan State University. We thank J. Martinez K. Doolittle, and B. Wurst for assistance on behavioral data collection and brain extractions, L. Racey, R. Mobley, J. Schuster, M. Tillotson, and E. Weigel for fish collection and/or fish care, and other members of the Boughman laboratory for additional support. A. Bell, H. Eisthen, J. Gallant, and M. Lucas gave valuable assistance while developing the brain extraction procedure. R. Harris extracted the RNA. D. Arasappan and B. Goetz gave bioinformatics training. B. Young gave bioinformatics training and extensive feedback on different versions of this manuscript. C. Weitecamp, C. Ofria, D. Bolnick, and M. Cummings gave other forms of support.

## Author Contributions

JK conceived of the study with input from HAH and JWB. JK, HAH, and JWB secured funding for the project and co-wrote the manuscript. JK collected the data and performed all bioinformatics and statistical analyses and generated the figures.

## References

1. Albert AYK, Millar NP, Schluter D. 2007. Character displacement of male nuptial colour in threespine sticklebacks (*Gasterosteus aculeatus*). Biol. J. Linn. Soc. 91:37–48.

2. Andersson M. 1994. Sexual Selection. Princeton University Press

3. Bloch NI, Corral-López A, Buechel SD, Kotrschal A, Kolm N, Mank JE. 2018. Early neurogenomic response associated with variation in guppy female mate preference. *Nat*. Ecol. Evol. 2:1772–1781.

4. Bolnick DI, Barrett RDH, Oke KB, Rennison DJ, Stuart YE. 2018. (Non)parallel evolution. Annu. Rev. Ecol. Evol. Syst. 49:303–330.

5. Boughman JW. 2001. Divergent sexual selection enhances reproductive isolation in sticklebacks. Nature 411:944–948.

6. Boughman JW. 2007. Condition-dependent expression of red colour differs between stickleback species. J. Evol. Biol. 20:1577–1590.

7. Boughman JW, Rundle HD, Schluter D. 2005. Parallel evolution of sexual isolation in sticklebacks. Evolution 59:361–373.

8. Conte GL, Arnegard ME, Best J, Chan YF, Jones FC, Kingsley DM, Schluter D, Peichel CL. 2015. Extent of QTL reuse during repeated phenotypic divergence of sympatric threespine stickleback. Genetics 201:1189–1200.

9. Conte GL, Schluter D. 2013. Experimental confirmation that body size determines mate preference via phenotype matching in a stickleback species pair. Evolution 67:1477–84.

10. Cummings ME, Larkins-Ford J, Reilly CRL, Wong RY, Ramsey M, Hofmann HA. 2008. Sexual and social stimuli elicit rapid and contrasting genomic responses. Proc. R. Soc. B Biol. Sci. 275:393–402.

11. DeAngelis RS, Hofmann HA. 2020. Neural and molecular mechanisms underlying female mate choice decisions in vertebrates. J. Exp. Biol. 223:jeb207324.

12. Endler JA, Houde AE. 1995. Geographic variation in female preferences and for male traits in *Poecilia reticulata*. Evolution 49:456–468.

13. Fox J, Weisberg S. 2019. An R Companion to Applied Regression. 3rd ed. Thousand Oaks CA: Sage Available from: https://socialsciences.mcmaster.ca/jfox/Books/Companion

14. Gerhardt HC, Brooks R. 2009. Experimental analysis of multivariate female choice in gray treefrogs (*Hyla versicolor*: Evidence for directional and stabilizing selection. Evolution 63:2504–2512.

15. Goodson J. 2008. Nonapeptides and the evolutionary patterning of sociality. In: Progress in Brain Research. Vol. 170. Elsevier. p. 3–15. Available from: https://linkinghub.elsevier.com/retrieve/pii/S0079612308004019

16. Gore A. 2002. GnRH: The Master Molecule of Reproduction. Springer

17. Gow JL, Peichel CL, Taylor EB. 2006. Contrasting hybridization rates between sympatric three-spined sticklebacks highlight the fragility of reproductive barriers between evolutionarily young species. Mol. Ecol. 15:739–752.

18. Härer A, Bolnick DI, Rennison DJ. 2021. The genomic signature of ecological divergence along the benthic-limnetic axis in allopatric and sympatric threespine stickleback. Mol. Ecol. 30:451–463.

19. Head ML, Kozak GM, Boughman JW. 2013. Female mate preferences for male body size and shape promote sexual isolation in threespine sticklebacks. Ecol. Evol. 3:2183–2196.

20. Jacobs A, Carruthers M, Yurchenko A, Gordeeva NV, Alekseyev SS, Hooker O, Leong JS, Minkley DR, Rondeau EB, Koop BF, et al. 2020. Parallelism in eco-morphology and gene expression despite variable evolutionary and genomic backgrounds in a Holarctic fish.B PLOS Genet. 16:e1008658.

21. Jennions MD, Petrie M. 2007. Variation in mate choice and mating preferences: A review of causes and consequences. Biol. Rev. 72:283–327.

22. Jones FC, Chan YF, Schmutz J, Grimwood J, Brady SD, Southwick AM, Absher DM, Myers RM, Reimchen TE, Deagle BE, et al. 2012. A genome-wide SNP genotyping array reveals patterns of global and repeated species-pair divergence in sticklebacks. Curr. Biol. 22:83–90.

23. Juntti SA, Hilliard AT, Kent KR, Kumar A, Nguyen A, Jimenez MA, Loveland JL, Mourrain P, Fernald RD. 2016. A neural basis for control of cichlid female reproductive behavior by Prostaglandin F2α. Curr. Biol. 26:943–949.

24. Keagy J, Lettieri L, Boughman JW. 2016. Male competition fitness landscapes predict both forward and reverse speciation. Ecol. Lett. 19:71–80.

25. Kidd MR, Dijkstra PD, Alcott C, Lavee D, Ma J, O’Connell LA, Hofmann HA. 2013. Prostaglandin F2α facilitates female mating behavior based on male performance. Behav. Ecol. Sociobiol. 67:1307– 1315.

26. Kolberg L, Raudvere U, Kuzmin I, Vilo J, Peterson H. 2020. gprofiler2 –– an R package for gene list functional enrichment analysis and namespace conversion toolset g:Profiler. F1000Research 9(ELIXIR):709.

27. Kozak GM, Reisland M, Boughmann JW. 2009. Sex differences in mate recognition and conspecific preference in species with mutual mate choice. Evolution 63:353–365.

28. Kraaijeveld K, Kraaijeveld-Smit FJL, Maan ME. 2011. Sexual selection and speciation: The comparative evidence revisited. Biol. Rev. 86:367–377.

29. Lackey A, Boughman J. 2013. Divergent sexual selection via male competition: ecology is key. J. Evol. Biol. 26:1611–1624.

30. Lackey ACR, Boughman JW. 2014. Female discrimination against heterospecific mates does not depend on mating habitat. Behav. Ecol. 25:1256–1267.

31. Langfelder P, Horvath S. 2008. WGCNA: An R package for weighted correlation network analysis. BMC Bioinformatics 9:559.

32. Langfelder P, Horvath S. 2012. Fast R functions for robust correlations and hierarchical clustering. J. Stat. Softw. 46:1–17.

33. Lerner TN, Holloway AL, Seiler JL. 2021. Dopamine, updated: Reward prediction error and beyond. Curr. Opin. Neurobiol. 67:123–130.

34. Love MI, Huber W, Anders S. 2014. Moderated estimation of fold change and dispersion for RNA-seq data with DESeq2. Genome Biol. 15:550.

35. Mani GS, Clarke BC. 1997. Mutational order: A major stochastic process in evolution. Proc. R. Soc. Lond. B Biol. Sci. 240:29–37.

36. Marvel M, Levavi-Sivan B, Wong T-T, Zmora N, Zohar Y. 2021. Gnrh2 maintains reproduction in fasting zebrafish through dynamic neuronal projection changes and regulation of gonadotropin synthesis, oogenesis, and reproductive behaviors. Sci. Rep. 11:6657.

37. McKinnon JS, Mori S, Blackman BK, David L, Kingsley DM, Jamieson L, Chou J, Schluter D. 2004. Evidence for ecology’s role in speciation. Nature 429:294–298.

38. McPhail J. 1994. Speciation and the evolution of reproductive isolation in the sticklebacks (*Gasterosteus*) of south-western British Columbia. In: Bell M, Foster S, editors. The Evolutionary Biology of the Threespine Stickleback. Oxford: Oxford Science Publications. p. 399–437.

39. Mendelson TC, Martin MD, Flaxman SM. 2014. Mutation-order divergence by sexual selection: Diversification of sexual signals in similar environments as a first step in speciation. Ecol. Lett. 17:1053–1066.

40. Mendelson TC, Safran RJ. 2021. Speciation by sexual selection: 20 years of progress. Trends Ecol. Evol. 36:1153–1163.

41. Mobley RB, Tillotson ML, Boughman JW. 2016. Olfactory perception of mates in ecologically divergent stickleback: population parallels and differences. Evol. Ecol. Res. 17:551–564.

42. Nagel L, Schluter D. 1998. Body size, natural selection, and speciation in sticklebacks. Evolution 52:209– 218.

43. Oh KP, Shaw KL. 2013. Multivariate sexual selection in a rapidly evolving speciation phenotype. Proc. R. Soc. B Biol. Sci. 280:20130482.

44. Okuyama T, Yokoi S, Abe H, Isoe Y, Suehiro Y, Imada H, Tanaka M, Kawasaki T, Yuba S, Taniguchi Y, et al. 2014. A neural mechanism underlying mating preferences for familiar individuals in medaka fish. Science 343:91–94.

45. Panhuis TM, Butlin R, Zuk M, Tregenza T. 2001. Sexual selection and speciation. Trends Ecol. Evol. 16:364–371.

46. Pfenning AR, Hara E, Whitney O, Rivas MV, Wang R, Roulhac PL, Howard JT, Wirthlin M, Lovell PV, Ganapathy G, et al. 2014. Convergent transcriptional specializations in the brains of humans and song-learning birds. Science 346:1256846.

47. R Core Team. 2022. R: A language and environment for statistical computing. R Foundation for Statistical Computing, Vienna, Austria. URL https://www.R-project.org/. Available from: https://www.R-project.org/

48. Renn SCP, Machado HE, Duftner N, Sessa AK, Harris RM, Hofmann HA. 2018. Gene expression signatures of mating system evolution. Genome 61:287–297.

49. Ritchie MG. 2007. Sexual selection and speciation. Annu. Rev. Ecol. Evol. Syst. 38:79–102.

50. Rosenthal GG, Ryan MJ. 2022. Sexual selection and the ascent of women: Mate choice research since Darwin. Science 375:eabi6308.

51. Rundle HD, Nagel L, Boughman JW, Schluter D. 2000. Natural selection and parallel speciation in sympatric sticklebacks. Science 287:306–308.

52. Rundle HD, Schluter D. 1998. Reinforcement of stickleback mate preferences: Sympatry breeds contempt. Evolution 52:200–208.

53. Ryan MJ. 2021. Darwin, sexual selection, and the brain. Proc. Natl. Acad. Sci. 118:e2008194118.

54. Ryan MJ, Rosenthal GG. 2001. Variation and selection in swordtails. In: Dugatkin L, editor. Model Systems in Behavioral Ecology. Princeton: Princeton University Press. p. 133–148.

55. Schluter D. 2009. Evidence for ecological speciation and Its alternative. Science 323:737–741.

56. Schluter D, McPhail JD. 1992. Ecological character displacement and speciation in sticklebacks. Am. Nat. 140:85–108.

57. Scordato ES, Safran RJ. 2014. Geographic variation in sexual selection and implications for speciation in the Barn Swallow. Avian Res. 5:8.

58. Servedio MR, Boughman JW. 2017. The role of sexual selection in local adaptation and speciation. Annu. Rev. Ecol. Evol. Syst. 48:85–109.

59. Sinha S, Jones BM, Traniello IM, Bukhari SA, Halfon MS, Hofmann HA, Huang S, Katz PS, Keagy J, Lynch VJ, et al. 2020. Behavior-related gene regulatory networks: A new level of organization in the brain. Proc. Natl. Acad. Sci. 117:23270–23279.

60. Soria-Carrasco V, Gompert Z, Comeault AA, Farkas TE, Parchman TL, Johnston JS, Buerkle CA, Feder JL, Bast J, Schwander T, et al. 2014. Stick insect genomes reveal natural selection’s role in parallel speciation. Science 344:738–742.

61. Storey JD, Tibshirani R. 2003. Statistical significance for genomewide studies. Proc. Natl. Acad. Sci. 100:9440–9445.

62. Taylor EB, McPhail J. 2000. Historical contingency and ecological determinism interact to prime speciation in sticklebacks, *Gasterosteus*. Proc. R. Soc. Lond. B Biol. Sci. 267:2375–2384.

63. Taylor EB, McPhail JD. 1999. Evolutionary history of an adaptive radiation in species pairs of threespine sticklebacks (*Gasterosteus*): Insights from mitochondrial DNA. Biol. J. Linn. Soc. 66:271–291.

64. Temple JL, Millar RP, Rissman EF. 2003. An evolutionarily conserved form of gonadotropin-releasing hormone coordinates energy and reproductive behavior. Endocrinology 144:13–19.

65. Tinghitella RM, Weigel EG, Head M, Boughman JW. 2013. Flexible mate choice when mates are rare and time is short. Ecol. Evol. 3:2820–2831.

66. Vamosi SM. 2002. Predation sharpens the adaptive peaks: Survival trade-offs in sympatric sticklebacks. Ann. Zool. Fenn. 39:237–248.

67. Weitekamp CA, Hofmann HA. 2017. Brain systems underlying social behavior. In: Evolution of Nervous Systems. 2nd ed. Elsevier. p. 327–334. Available from: 10.1016/B978-0-12-804042-3.00025-7

68. Wong RY, Ramsey ME, Cummings ME. 2012. Localizing brain regions associated with female mate preference behavior in a swordtail. PLoS ONE 7:e50355.

69. Young RL, Ferkin MH, Ockendon-Powell NF, Orr VN, Phelps SM, Pogány Á, Richards-Zawacki CL, Summers K, Székely T, Trainor BC, et al. 2019. Conserved transcriptomic profiles underpin monogamy across vertebrates. Proc. Natl. Acad. Sci. 116:1331–1336.

70. Young RL, Weitekamp CA, Triki Z, Su Y, Bshary R, Hofmann HA. 2022. Shared neural transcriptomic patterns underlie the repeated evolution of mutualistic cleaning behavior in Labridae wrasses. EcoEvoRxiv preprint.

