## Supplementary Material for "Mate choice in the brain: Species differ in how male traits ‘turn on’ gene expression in female brains"

**This document includes:**

Supplementary Materials and Methods
Supplementary Figures 1 to 4
Legends for Supplementary Tables 1 to 10
Supplementary References

### Supplementary Materials and Methods

*Enticing males to nest:* Reproductive males (based on nuptial coloration) were taken from holding tanks and placed individually in visually isolated 110-L aquaria (with a 76 x 31 cm footprint and water depth of 43 cm) with nesting materials (plastic tray of sand, an inverted half of a terra cotta flowerpot for cover, a plastic plant, and filamentous algae (*Chara* sp.)). To entice males to build a nest and perform courtship behaviors, we presented them with a gravid conspecific female from their native lake in a glass jar with screen over the mouth once a day for five minutes. Males were given 14 days to nest; if they did not build a nest by that time they were removed from the experiment, the tank was cleaned, and all nesting materials replaced.

*Additional details about behavioral trials:* All females were weighed and photographed under standardized conditions prior to behavioral trials (Keagy et al. 2016). We used the data from the courtship trials to construct a preference score on a 0-4 point scale with 1-point increments indicating increased preference. A score of 0 corresponds to no behaviors indicating interest in the male and a score of 4 corresponds to laying eggs in the nest (Kozak et al. 2009; Kozak and Boughman 2009; Lackey and Boughman 2013; Tinghitella et al. 2013). We averaged the preference score for the two observers (there were 3 cases of minor disagreement out of 57) and then analyzed this data using an ANOVA with type-II sums of squares with lake, species, treatment, and all possible interactions as fixed effects (Fig. 1B). In prior work, we found that after 20 min together, males became very aggressive to females if they did not mate; therefore, we typically end courtship trials at 20 min (Boughman 2001; Kozak et al. 2009; Kozak and Boughman 2009; Lackey and Boughman 2013; Tinghitella et al. 2013). Spawning was uncommon, with 9/57 trials (16%) in the complete behavioral dataset, or 6/53 (11%) for the gene expression dataset. Courtship trials were immediately ended when spawning occurred because afterwards males become very aggressive to females (Kozak et al. 2009; Kozak and Boughman 2009; Lackey and Boughman 2013; Tinghitella et al. 2013). The duration of trials where spawning occurred was 203 – 1200 s (3.38 – 20 min).

*Male morphological data:* Each male was used only once in a courtship trial, after which point it was marked with an elastomer tag (Northwest Marine Technology, Shaw Island, WA) for future identification. Using a 0-5 scale with 0.5 increments, we then assessed 1) the area of the throat that was red, 2) the intensity of this red coloration, 3) the intensity of the blue eye coloration, 4) the intensity of the body coloration, and 5) the darkness of the body (Boughman 2001; Boughman 2007; Lewandowski and Boughman 2008; Keagy et al. 2016). The following afternoon, the same males were photographed and weighed under standardized conditions. Photographs were used to determine standard length - the distance from the tip of the snout to the caudal peduncle (Keagy et al. 2016).

*Description of PCs:* Three principal component (PC) analyses were conducted to describe variation in male morphology, male behavior, and female behavior. Female behavioral traits were 1) the proportion of times males led females to the nest and she followed (a measure related to interest in courtship (Kozak et al. 2009)), 2) the proportion of times males showed the nest to a female and she entered (a measure related to interest in mating(Kozak et al. 2009)), and 3) the preference score described above (Supplementary Fig. 1A). Male behavioral traits were 1) the number of approaches to the female, 2) the number of zig-zag dances, 3) the number of leads to the nest, 4) the number of bites, and 5) the number of chases (Supplementary Fig. 1B). Male morphological traits were 1) a throat score calculated by summing the score of the area of throat coloration and the score of the intensity of this throat coloration (Lewandowski and Boughman 2008; Keagy et al. 2016), 2) eye color intensity, 3) body color intensity, 4) body darkness, and 5) standard length, a common measure of size (Supplementary Fig. 1C). Traits were standardized prior to principal components analysis, and for male behavioral traits, log-transformed prior to standardization.

Fbehav-PC1 represented female responsiveness (explaining 83% of the total trait variation) and differentiated females that showed consistently low versus high preferences for the males courting them. Fbehav-PC2 differentiated females that primarily displayed behaviors indicating interest early in courtship versus later in courtship (explaining 13% of the total variation). Mbehav-PC1 (54%) described courtship vigor, with higher scores indicating fish that did more courtship behaviors. Mbehav-PC2 (29%) described the strategy employed by males in courtship, differentiating fish that used display elements such as zig-zag dance and leads to nest from fish that used aggressive elements such as biting and chasing. This is a well-described difference between limnetic and benthic fish respectively, although there is plasticity in this behavior (Nagel and Schluter 1998; Kozak et al. 2009). Mmorph-PC1 (40%) differentiated large and dull (typically benthic) fish from small and colorful (typically limnetic) fish. Mmorph-PC2 (25%) differentiated large fish with dark and intensely colored bodies from small fish with no body coloration.

### Supplementary Figures

A *Female Mate Preference*

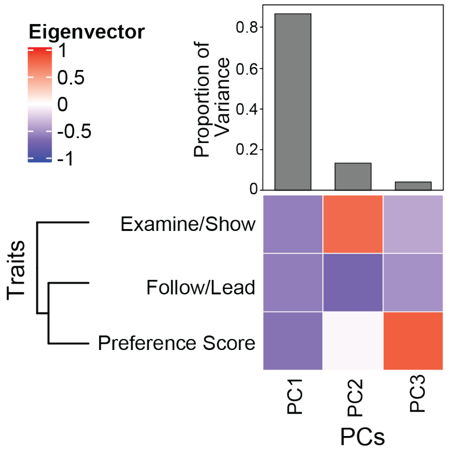


| | PC1 | PC2 | PC3 |
|---|---|---|---|
| **Proportion of Variance** | 0.83 | 0.13 | 0.04 |
| **Cumulative Variance** | 0.83 | 0.96 | 1.00 |
| **Eigenvalues** | 2.50 | 0.38 | 0.12 |

B *Male behavior*

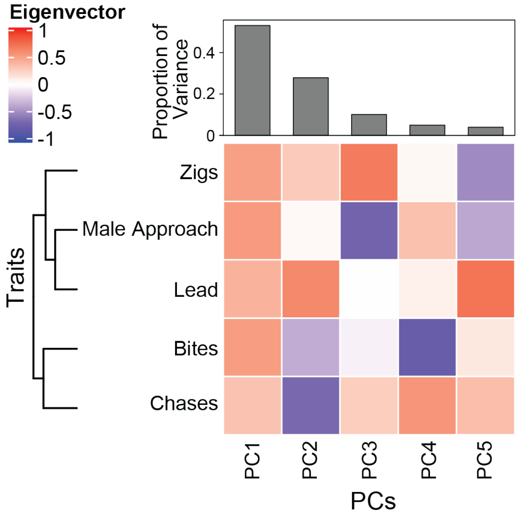


| | PC1 | PC2 | PC3 | PC4 | PC5 |
|---|---|---|---|---|---|
| **Proportion of Variance** | 0.53 | 0.29 | 0.10 | 0.05 | 0.04 |
| **Cumulative Variance** | 0.53 | 0.81 | 0.91 | 0.96 | 1.00 |
| **Eigenvalues** | 2.66 | 1.39 | 0.51 | 0.25 | 0.20 |

C *Male morphology*

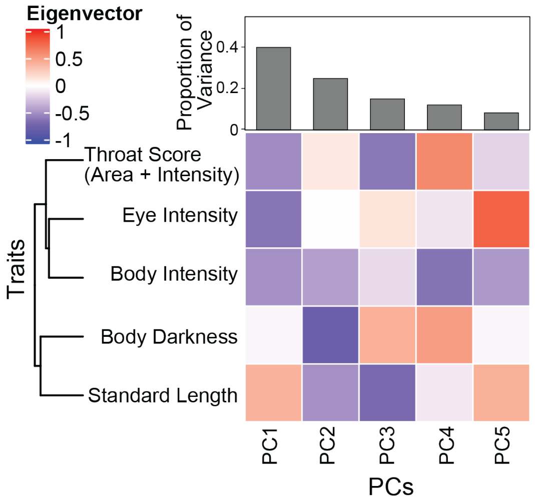


| | PC1 | PC2 | PC3 | PC4 | PC5 |
|---|---|---|---|---|---|
| **Proportion of Variance** | 0.40 | 0.25 | 0.15 | 0.12 | 0.08 |
| **Cumulative Variance** | 0.40 | 0.65 | 0.80 | 0.92 | 1.00 |
| **Eigenvalues** | 1.99 | 1.24 | 0.75 | 0.60 | 0.41 |

**Supplementary Figure 1. Principal Components Analysis of (A) Female mate preference behaviors, (B) Male courtship behaviors, and (C) Male sexually selected morphological characters.** Figures are visual representation of information contained in tables. The first two principal components from each of these analyses were used in further analyses. See "Supplemental Materials and Methods" for definitions of the traits included in the PCAs.

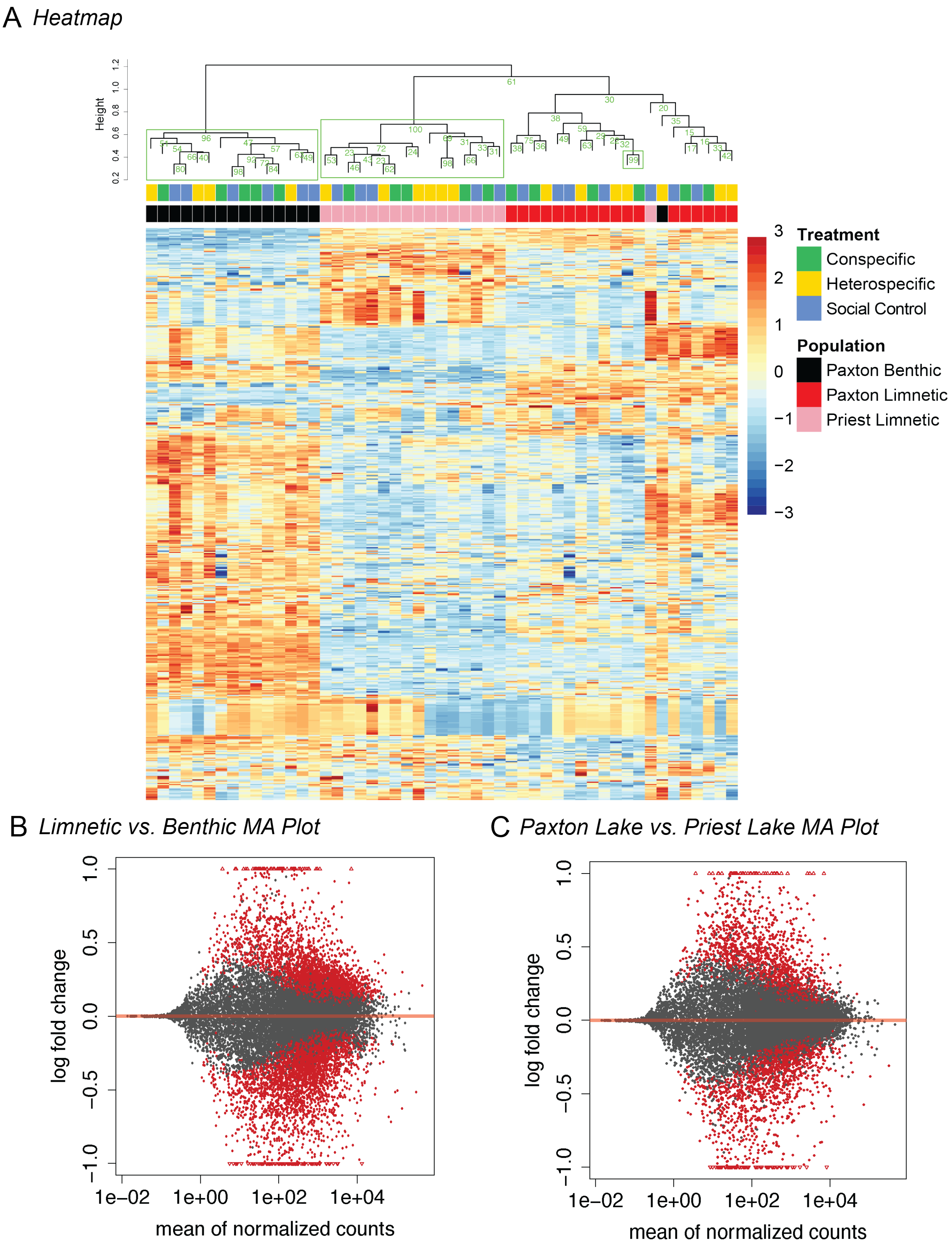
A Heatmap
Height
1.2
1.0
0.8
0.6
0.4
0.2
Treatment
Conspecific
Heterospecific
Social Control
Population
Paxton Benthic
Paxton Limnetic
Priest Limnetic
3
2
1
0
−1
−2
−3
B Limnetic vs. Benthic MA Plot
C Paxton Lake vs. Priest Lake MA Plot
log fold change
1.0
0.5
0.0
−0.5
−1.0
1e−02
1e+00
1e+02
1e+04
mean of normalized counts

**Supplementary Figure 2. Population differences in gene expression. (A) Heatmap of z-scores of variance-stabilized normalized counts for 500 most variable genes across all populations.** Rows are genes and columns are individual brains. Dendrogram of individual samples has bootstrap p-values in green. Green rectangles indicate groupings with >95% bootstrap support. **(B) MA plots of differential gene expression between limnetic and benthic fish from Paxton Lake and (C) limnetic fish from Paxton and Priest Lakes.** Genes that are significantly differentially expressed are in red. We used a significance threshold of FDR corrected $P < 0.1$. Triangles indicate points which are outside the plot window (i.e., log2-fold change > 1).

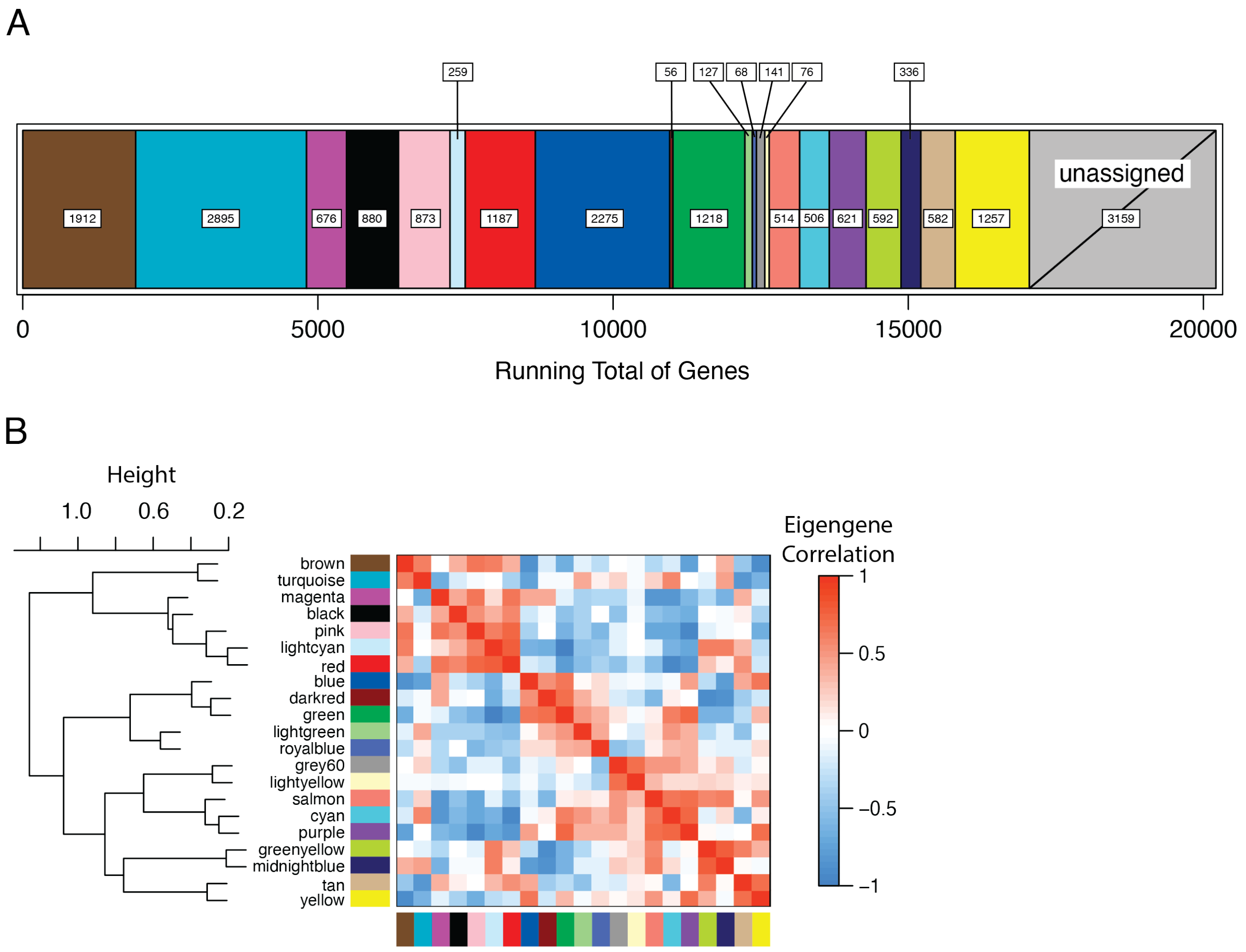


**Supplementary Figure 3. WGCNA modules. (A) Relative size of each module.** Number of genes in white boxes. **(B) Module relationships.** On the left is the dendrogram with shorter distances between nodes indicating closer relationships. The correlogram shows blocks of similar modules in red.

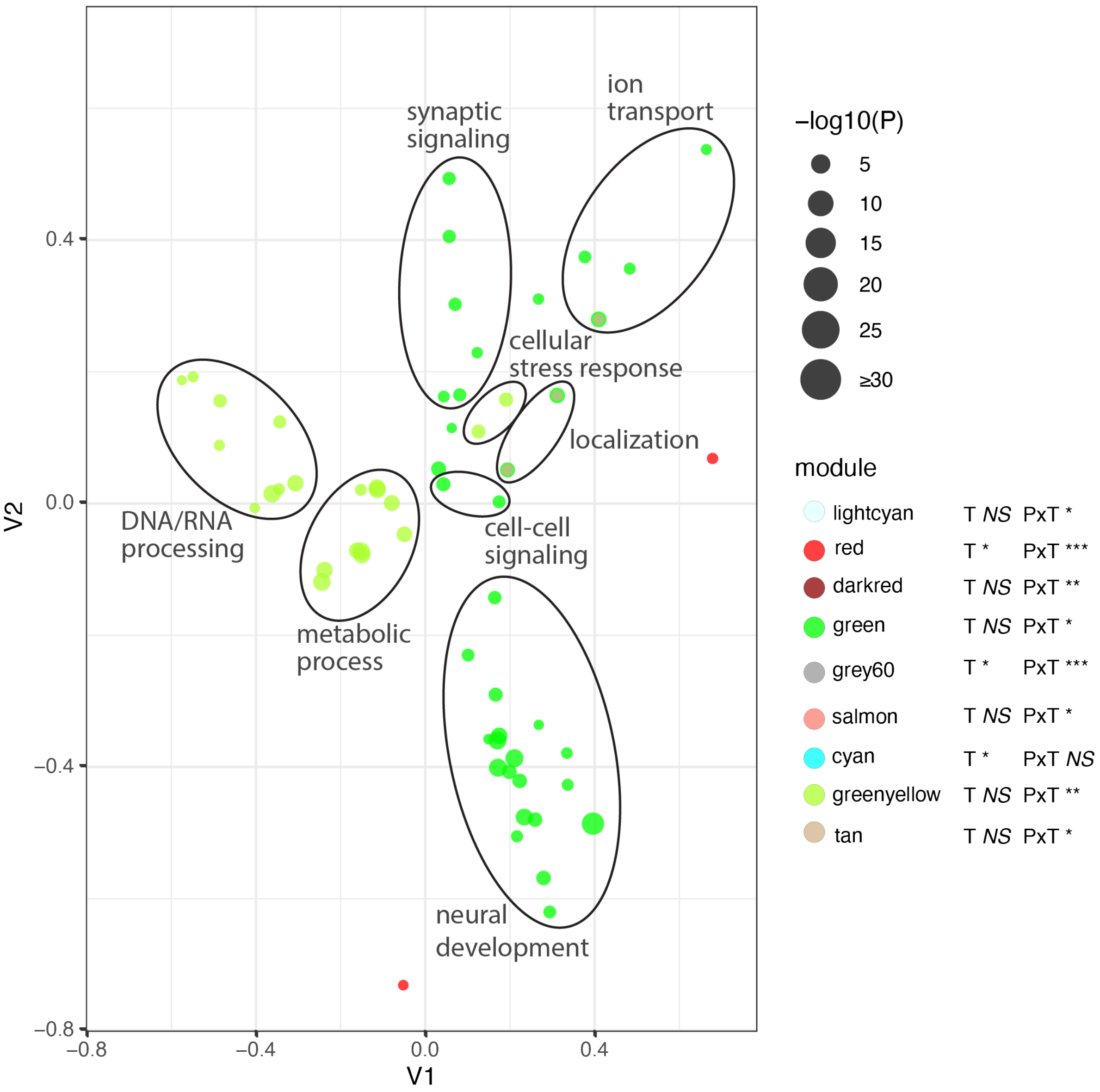


**Supplementary Figure 4. Biological Processes GO terms for modules that had population by female behavior PC2 interactions.** Eight modules had significant population by Fbehav-PC2 interactions. Statistical significance of the main effect of Fbehav-PC2 and the population by Fbehav-PC2 interaction effect are indicated as: *** < 0.001, ** < 0.01, * < 0.05. GO terms for genes in these modules that were overrepresented (significance indicated by circle size) were first quantified for semantic similarity and then plotted as MDS graphs. Groups of terms with similar functions are circled and manually annotated to aid interpretation.

**Legends for Supplementary Tables 1 to 10**

Supplementary tables are contained as separate sheets in an Excel workbook called "Supplementary Tables.xlsx"

**Supplementary Table 1.** Raw gene expression count data.

**Supplementary Table 2.** Results of statistical tests for differential expression of genes comparing limnetic vs. benthic fish in Paxton Lake and limnetic fish from Paxton vs. Priest Lakes

**Supplementary Table 3.** Results of statistical tests for differential expression of genes comparing different treatments, separately for each population.

**Supplementary Table 4.** GO analysis for differentially expressed genes (DEGs) identified in Supplementary Table 3.

**Supplementary Table 5.** WGCNA eigengene expression table.

**Supplementary Table 6.** Results of statistical tests (t-tests) for differential expression of module eigengenes for comparisons between specific populations and treatments.

**Supplementary Table 7.** WGCNA linear model significance values for population, traits, and trait by population interactions.

**Supplementary Table 8.** WGCNA GO analysis for each module.

**Supplementary Table 9.** Candidate gene linear model significance values for population, traits, and trait by population interactions.

**Supplementary Table 10.** Variance-stabilized normalized counts; these form the basis of the WGCNA and candidate gene analyses.

### Supplementary References


Boughman JW. 2001. Divergent sexual selection enhances reproductive isolation in sticklebacks. *Nature* 411:944–948.

Boughman JW. 2007. Condition-dependent expression of red colour differs between stickleback species. *J. Evol. Biol.* 20:1577–1590.

Keagy J, Lettieri L, Boughman JW. 2016. Male competition fitness landscapes predict both forward and reverse speciation. *Ecol. Lett.* 19:71–80.

Kozak GM, Boughman JW. 2009. Learned conspecific mate preference in a species pair of sticklebacks. *Behav. Ecol.* 20:1282–1288.

Kozak GM, Reisland M, Boughmann JW. 2009. Sex differences in mate recognition and conspecific preference in species with mutual mate choice. *Evolution* 63:353–365.

Lackey ACR, Boughman JW. 2013. Loss of sexual isolation in a hybridizing stickleback species pair. *Curr. Zool.* 59:591–603.

Lewandowski E, Boughman J. 2008. Effects of genetics and light environment on colour expression in threespine sticklebacks. *Biol. J. Linn. Soc.* 94:663–673.

Nagel L, Schluter D. 1998. Body size, natural selection, and speciation in sticklebacks. *Evolution* 52:209–218.

Tinghitella RM, Weigel EG, Head M, Boughman JW. 2013. Flexible mate choice when mates are rare and time is short. *Ecol. Evol.* 3:2820–2831.
